# Metabolic Rigidity as a Mechanical Barrier to Malaria: Flickering Loss in PKLR-Deficient Erythrocytes

**DOI:** 10.1101/2025.07.29.667448

**Authors:** Natalia Hernando-Ospina, Macarena Calero, Guillermo Solís, Isabel G. Azcárate, Diego Herráez-Aguilar, Niccolò Caselli, Lara H. Moleiro, Jose-Carlos Segovia, José M. Bautista, Francisco Monroy

## Abstract

Pyruvate kinase (PK) deficiency is a rare hereditary enzymopathy caused by mutations in the *PKLR* gene, leading to reduced glycolytic ATP production in red blood cells (RBCs) and contributing to chronic hemolytic anemia. Here, we use high-speed flickering spectroscopy and passive microrheology to assess how ATP depletion reshapes the nanomechanical properties of RBC membranes. Compared to healthy controls, *PKLR*-mutant erythrocytes exhibit marked reductions in ATP-dependent flickering amplitude and membrane fluidity, consistent with impaired metabolic elasticity. Strikingly, when infected with *Plasmodium yoelii*, these metabolically rigidified cells retain mechanical properties that appear to hinder parasite-induced membrane remodeling. By mapping single-cell viscoelastic landscapes across healthy, mutated, infected and coinfected mouse RBC populations, we uncover a potential biomechanical barrier against malaria imposed by glycolytic insufficiency. These findings highlight a mechanobiological axis of host resistance and position label-free flickering analysis as a powerful tool for diagnosing RBC enzymopathies and probing infection susceptibility at the single-cell level.

## INTRODUCTION

Red blood cells (RBCs), or erythrocytes, are highly specialized, anucleate cells uniquely adapted to transport respiratory gases throughout the circulatory system [1,2]. Lacking mitochondria and other organelles, mature RBCs rely exclusively on anaerobic glycolysis to meet their energy demands [1,3]. Central to this pathway is the Embden–Meyerhof (EM) process [4], whose final step is catalyzed by pyruvate kinase (PK)—a transphosphorylating enzyme encoded by the *PKLR* gene in erythroid precursors [1–4]. Specifically, PK irreversibly converts phosphoenolpyruvate into pyruvate, generating one of the two net ATP molecules that support RBC homeostasis [3,4]. This limited ATP pool is essential for maintaining ion gradients, redox balance, and, critically, membrane deformability. The remarkable elasticity of RBCs, required for their passage through capillaries narrower than their resting diameter, arises from an intricate membrane architecture composed of a spectrin-based cytoskeleton anchored to a highly structured lipid bilayer [5]. Together, these components form a dynamic, mechanoelastic shell that sustains repeated deformation while preserving cell integrity and function in circulation [1–5]. Mutations in the *PKLR* gene impair PK function, leading to inherited pyruvate kinase deficiency (PKD), one of the most common causes of congenital nonspherocytic hemolytic anemia [6, 7]. In homozygous individuals, insufficient ATP production triggers ion leakage, dysregulated membrane potential, and cellular dehydration, ultimately compromising the structural resilience of RBCs with no compensatory gene expression to offset this defect [7]. These altered mechanics promote premature removal from the bloodstream due to reduced deformability and increased susceptibility to hemolysis [8]. *PKLR*-linked enzymopathies have been documented worldwide, though they are most prevalent in individuals of Northern European ancestry [9], raising the hypothesis that PKD may confer a selective advantage [9, 10]. Interestingly, several epidemiological and functional studies suggest that PKD may confer partial protection against malaria [10, 11], similar to the protective effects observed with other RBC polymorphisms, such as those in the PIEZO1, G6PD, or HBB genes [12,13]. This PKD protective effect is thought to arise from the reduced metabolic plasticity and increased membrane stiffness of PKD-RBCs, which may interfere with *Plasmodium* spp. invasion, maturation or egress. Because successful intraerythrocytic development of the parasite requires extensive remodelling of the host cytoskeleton and sustained energy availability, metabolically compromised RBCs may create a biomechanically hostile environment for parasite survival.

Functionally, PKD-RBCs fail to maintain the membrane elasticity required for healthy circulation [6, 7]. Potassium and water leak out, while intracellular calcium and sodium levels increase, further altering the mechanical and osmotic balance of the RBC membrane [14,15]. In pathophysiological terms, this biophysical imbalance contributes to a more rigid cell morphology and loss of flexibility, rendering the cells vulnerable to shear forces and splenic clearance [16]. Given the strict dependence of RBC membrane flexibility on glycolytic ATP, we hypothesized that genetic ATP depletion, such as that seen in PKLR deficiency, may alter erythrocyte biomechanics in a manner that interferes with intracellular parasite development in malaria infection [17]. Live-cell imaging has revealed that healthy RBCs display dynamic membrane undulations—termed “flickering” [18], which reflect an active, energy-dependent mechanical phenotype indicative of cellular fitness. Initially attributed to thermal fluctuations [19], flickering is now widely recognized as a metabolically active process [20,21,22,23,24,25]. High-speed microscopy has revealed that active flickering arises from spectrin network remodelling [26,27], and ATP-dependent shape deformations mediated by a cytoskeleton-membrane nonequilibrium dynamics [28,29]. In contrast, ATP-depleted RBCs exhibit markedly reduced flickering, indicative of a passivated mechanical state [23,30,31], supporting the use of observable flickering as a functional readout of membrane responsiveness [32,33]. In pathological contexts, such as parasitic RBC invasion, membrane flickering reflects the altered energy landscape of the membrane–cytoskeleton composite and thus serves as a proxy for the mechanical readiness of the cell to undergo deformation events such as Plasmodium invasion.

A growing body of translational research underscores the pivotal role of RBC biomechanics in modulating susceptibility to malaria infection [16,17,34]. Successful invasion following intraerythrocytic development of *Plasmodium* spp. depend on extensive remodelling of the host RBC membrane and the insertion of parasite-derived proteins that alter the mechanical properties of the cell [35,36]. Accordingly, the biomechanical plasticity of the RBC membrane is a critical determinant of parasite fitness, invasion efficiency, and replication success [37]. Notably, suppression of ATP-dependent membrane flickering in metabolically compromised RBCs has been associated with elevated membrane tension [38,39], which increases the energy barrier for merozoite-induced membrane wrapping [40]. Since membrane tension has been identified as a mechanical barrier to invasion [37,38], diminished flickering amplitudes may serve as a biomechanical signature of partial resistance to malaria. Complementary theoretical models further support this notion [40,41], demonstrating that cytoskeletal resistance and membrane tension jointly govern the energetics of membrane wrapping during parasite entry. Membrane flickering spectroscopy thus emerges as a non-invasive method to infer key mechanical parameters at single-cell resolution [37]. Despite these important advances, the impact of genetically impaired RBCs—particularly those with reduced ATP availability due to metabolic enzymopathies such as *PKLR* deficiency—on malaria pathogenesis remains poorly understood. By resolving the viscoelastic flickering phenotype at the single-cell level, we aimed to determine whether *PKLR*-associated metabolic impairment establishes a mechanically rigid cellular environment that hinders parasite entry, survival, or replication.

In this work, we examined how *PKLR*-linked metabolic deficiency affects the mechanical phenotype of red blood cells and its potential role in modulating susceptibility to malaria. To address this gap, we employed a preclinical murine model to perform a comparative nanomechanical analysis of RBCs from *PKLR*-deficient and wild-type mice, both in uninfected states and following experimental infection with *Plasmodium yoelii*, a rodent malaria species widely used as an animal model for human lethal malaria [42]. Using high-speed flickering spectroscopy [27] in combination with passive microrheology [43], we quantified membrane dynamics, such as fluctuation amplitudes and rates, and viscoelastic properties e.g., stiffness and fluidity across individual cells. This integrative approach allowed us to dissect the independent and combined effects of genetic ATP depletion and parasitic infection on RBC deformability. *PKLR*-deficient RBCs exhibited significantly diminished flickering activity and reduced membrane fluidity, consistent with their impaired ATP production. Upon infection, these mechanically stiffened cells appeared less permissive to parasite-induced remodeling, supporting the idea that ATP-depletion establishes a biomechanical constraint on intracellular parasite development [44]. Within this framework, membrane flickering emerges as more than a passive reflection of cellular metabolism—it may function as a dynamic mechanical phenotype that contributes to host defence by limiting parasite invasion or maturation. Our findings suggest that reduced flickering, typically associated with pathological rigidity, could paradoxically represent a protective trait in the context of malaria infection. This positions flickering-based nanomechanical profiling as a promising approach for probing host-pathogen interactions and for identifying mechanical biomarkers of disease susceptibility in hereditary RBC disorders. Future studies extending this methodology to human samples and diverse parasitic stages may reveal new opportunities for diagnostic innovation and therapeutic targeting in malaria and other erythrocyte-related pathologies.

## MATERIALS AND METHODS

### Chemicals and buffers

Unless otherwise stated, all reagents and chemicals used in this study were obtained from Merck-Sigma-Aldrich (UK) and were of analytical grade. Deionized water was produced using a Milli-Q purification system (Millipore, USA) and used for all buffer preparations and dilutions. All solutions were freshly prepared, and pH was carefully adjusted and verified using a calibrated pH meter (Crison, Spain. Buffers used in RBC isolation and storage included phosphate-buffered saline (PBS), supplemented with glucose and bovine serum albumin (BSA). This PBS+ buffer was composed of 130 mM NaCl, 20 mM Na₃PO₄, 10 mM glucose, and 1 mg/mL BSA, adjusted to pH 7.4. For experiments requiring metabolic inhibition, a glucose-free variant of this buffer (PBS–) was prepared using the same ionic composition but omitting glucose. ATP-depleting agents included iodoacetamide (18 mM), used to alkylate thiol groups and inhibit glycolytic enzymes such as glyceraldehyde-3-phosphate dehydrogenase (G3PDH) and hexokinase (HK), and inosine (30 mM), which depletes intracellular phosphate and inhibits ATP regeneration. For cell fixation, glutaraldehyde (3% in PBS+) was used to cross-link membrane and cytoskeletal proteins and solidify intracellular hemoglobin. All stock solutions were prepared in PBS or Milli-Q water, filtered through 0.22 µm membranes where necessary, and stored at 4 °C for short-term use.

### Ethics statement

All animal experiments were approved by the Committee of Animal Experimentation of the Universidad Complutense de Madrid and conducted in accordance with Spanish (R.D. 53/2013) and European (Directive 2010/63/EU) regulations for the protection of animals used for scientific purposes. The procedures were performed simultaneously and under the same approved protocol as those previously reported [45].

### Mouse models for erythrocyte phenotyping

RBCs were isolated from two mouse genotypes: (i), PKLR-deficient mice generated by sets of recombinant inbred strains, congenic strains, and recombinant congenic strains identifying disruption of the *PKLR* gene, and (ii) wild-type C57BL/6J mice used as controls. Both strains are on a C57BL/6J genetic background and were previously described and characterized in detail [46,47]. RBCs were collected from three animals per group (n = 3 biological replicates per genotype). Blood was obtained by venipuncture from the caudal vein under brief isoflurane anesthesia. After collection, RBCs were immediately diluted in PBS+ buffer to preserve metabolic activity under physiological conditions.

### Experimental infection with *Plasmodium yoelii*

The rodent malaria parasite *Plasmodium yoelii yoelii* 17XL (PyL) was used for all infection experiments following the protocols previously described [48,49]. The parasite line was maintained in liquid nitrogen and revived from frozen stocks containing infected red blood cells (iRBCs) previously passaged in mice. Wild-type and PKLR-deficient C57BL/6J female mice, aged 8 weeks, were housed under standard conditions at the Animal Facility of the Universidad Complutense de Madrid. Environmental parameters included a 12:12 h light–dark cycle, controlled temperature (22–24 °C) and humidity (∼50%), and access to food (Teklad Global 18% Protein Rodent Diet, Harlan Laboratories) and water ad libitum. On day 0, mice were infected intraperitoneally with 2 × 10⁷ iRBCs from frozen PyL stocks. Parasitemia was monitored daily starting on day 2 post-infection by preparing thin blood smears stained with Wright’s eosin methylene blue. Parasite load was quantified microscopically. Mice were sacrificed at day 3 or day 6 post-infection based on parasitemia levels, and blood was collected for downstream analysis. Three independent mice were infected and analyzed per condition.

### Isolation and preparation of RBCs

Murine RBCs were isolated from freshly collected whole blood obtained by venipuncture under light anesthesia, in accordance with institutional animal care guidelines. Following collection, blood was diluted in phosphate-buffered saline supplemented with glucose and bovine serum albumin (PBS⁺), and RBCs were separated by centrifugation at 5,000 rpm for 10 minutes. The washing procedure was repeated three times to remove plasma and leukocytes. After the final wash, the RBC pellet was resuspended in PBS⁺ and stored at 37 °C for subsequent experiments.For each experimental condition, a total of 20 µL of whole blood was diluted into 1 mL of PBS+ buffer— phosphate-buffered saline supplemented with glucose and bovine serum albumin (BSA)—to maintain metabolic activity and osmotic balance. RBCs were purified by three sequential centrifugation steps at 5000 rpm for 10 minutes each at room temperature. After each centrifugation, the supernatant was carefully removed, and the pellet was resuspended in 1 mL of freshly prepared PBS⁺ to wash out plasma components and non-RBC contaminants. Following the third wash, the final RBC pellet was diluted 1:150 in PBS⁺ for microscopy and flickering spectroscopy assays and stored at 37 °C in a humidified incubator to mimic physiological conditions. This preparation protocol preserves the functional integrity and biophysical properties of the RBC membrane, including its dynamic flickering behavior [26], and has been previously validated in live-cell experiments combining high-speed imaging and fluctuation analysis of the membrane contour [27].

### RBC drug treatment

To investigate the metabolic and morphological responses of modified RBCs under glycolytic inhibition, specific metabolic modulators were used to alter ATP production. Iodoacetamide (18 mM), an alkylating agent that irreversibly inhibits glycolytic enzymes by targeting –SH groups on active sites (notably G3PDH and hexokinase), was combined with inosine (30 mM), a purine nucleoside that enhances ATP depletion by depleting intracellular phosphate reserves and preventing ATP regeneration via the G3PDH step. Both compounds were dissolved in PBS⁻ (phosphate-buffered saline lacking glucose), and 1 mL of this solution was added to the RBC pellet previously prepared and washed as described above. The cell suspension was incubated at 37 °C for 3 hours to ensure sufficient inhibition of glycolytic activity. Under these conditions, ATP levels are drastically reduced, and the phosphorylation of cytoskeletal and membrane-associated enzymes is effectively suppressed, leading to observable changes in membrane mechanics and flickering dynamics.

### Fixed RBC

To prepare fixed RBCs for reference measurements under fully arrested metabolic and mechanical conditions, chemical fixation was performed using glutaraldehyde. This cross-linking agent covalently binds amino groups in membrane and cytoskeletal proteins, leading to irreversible rigidification of the erythrocyte membrane and structural stabilization of hemoglobin. Following the final washing step, RBCs were resuspended in 1 mL of PBS⁺ and treated with 3% glutaraldehyde. The suspension was incubated for 2 hours at 37 °C to ensure complete fixation. This treatment results in ATP depletion, inhibition of membrane protein mobility, and the suppression of flickering dynamics, providing a passive mechanical baseline for comparison with metabolically active cells.

### Flickering spectroscopy

Fluctuation spectra of RBCs were acquired using high-speed phase-contrast video microscopy in frontal view [27]. Recordings were performed on a Nikon Eclipse Ti-2000 inverted microscope (Nikon, Japan) equipped with a 100 W TI-DH Pillar illuminator (TI-PS100W power supply), focused through a 0.72 LWD collimator for Köhler illumination under phase-contrast conditions. A high numerical aperture Plan Apo VC 100× oil immersion objective (NA 1.4) was used for optimal resolution. Time-resolved imaging was achieved using a FASTCAM SA3 ultrafast CMOS camera (Photron, San Diego), coupled via a variable optical magnifier (up to 2.25×) and the internal 1.5× magnification of the microscope base, resulting in a total magnification of 337.5×. This configuration yielded an effective pixel size of 50 × 50 nm² at the sample plane. Image sequences were recorded at 2000 frames per second (fps), with 5 s acquisition windows (10,000 frames per video). All measurements were performed at 37 °C to preserve physiological flickering activity.

### Flickering Data Analysis

The equatorial contour of each RBC was segmented from high-speed phase-contrast recordings using a custom tracking algorithm [27], implemented in a *Wolfram Mathematica* pipeline [32]. Each membrane profile was discretized into 1,024 equidistant points along the circumference of length 𝐿 = 𝜋𝐷 (where 𝐷 ≈ 6 − 8 𝜇𝑚 is the RBC diameter). the RBC diameter. For each tracked point—corresponding to a membrane element of characteristic size 𝑎 = 𝐿⁄1024 ≈ 10 𝑛𝑚 —a time series of radial displacements 𝑢_𝑘_(𝑡) was extracted over 5 s of acquisition at 2,000 frames per second, yielding high-resolution spatiotemporal datasets. These local membrane fluctuations were interpreted as the Brownian motion of submicron-sized membrane segments immersed in a viscoelastic environment characterized by a complex shear modulus *G̃* = *G^′^* + *iG^″^*, where *G^′^* is the storage shear modulus (elastic response) and *G^″^* = 𝜔𝜂 is the loss modulus related to the shear viscosity 𝜂 (viscous response). To extract these parameters, we employed the Mason–Weitz microrheology formalism [43], which relates the observed fluctuations to the complex frequency-dependent friction coefficient 𝜁̃(𝜔) ≡ 6𝜋𝑎𝜂̃, with related complex viscosity:

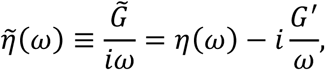

which incorporates both real viscous and imaginary elastic contributions. Hence, the complex diffusivity is then given by the generalized Stokes–Einstein relation:

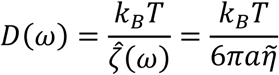

Here, 𝐷(𝜔) is the frequency-dependent diffusivity, 𝑎 is the effective hydrodynamic size of the fluctuating membrane element, and 𝜂̃ is the effectively complex viscosity of the surrounding medium (including viscoelastic contributions). The time-domain mean square displacement (MSD) was computed from the variance of the displacement trajectories, via the Einstein relation:

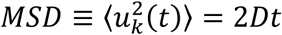

which is transformed to the frequency domain by Fourier-transform to extract the complex viscosity as [43]:

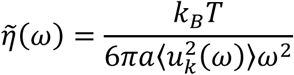

From this expression, we estimated local viscoelastic moduli: at high frequencies, the instantaneous elastic modulus was obtained as:

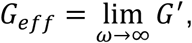

while at low frequencies, effective fluidity was defined as:

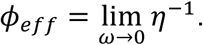

This microrheological approach allows label-free, spatially resolved quantification of the mechanical environment of RBC membranes, distinguishing passive thermal fluctuations in fixed or ATP-depleted cells from active flickering powered by metabolic energy [26,32].

### Statistics

To achieve statistical power in single-cell analyses, measurements from all three congenic animals per genotype were aggregated, allowing us to capture pooled phenotypic trends while controlling for intra-group variability. The statistical analyses were performed using *GraphPad Prism*, with graph visualization supported by *OriginLab*. Differences between experimental groups were assessed using one-way analysis of variance (ANOVA), with statistical significance defined according to standard thresholds (*p* < 0.05, unless otherwise stated). While our single-cell analyses uncovered statistically significant mechanical differences across conditions, the aggregation of data from three congenic mice per genotype necessitates caution in interpreting inter-individual variability. Preliminary variance decomposition suggests that within-group (intra-animal) variability dominates over between-animal (inter-animal) effects, justifying pooled analysis for detecting robust phenotypic trends.

## RESULTS

### ATP-Dependent Flickering Reveals Mechanical Signatures of Red Blood Cell Metabolism

We applied high-speed flickering spectroscopy to characterize the dynamic mechanical properties of RBC membranes from PKLR-deficient C57BL/6J mice under different metabolic states (see Methods). This non-invasive optical approach enables precise tracking of equatorial membrane fluctuations at nanometer resolution and kilohertz temporal sampling, yielding high-content biophysical readouts of live RBC mechanical behavior [26,32]. As shown in Figure 1A, the membrane contours of healthy RBCs (top) and glutaraldehyde-fixed RBCs (bottom) were segmented from phase-contrast micrographs and reconstructed with subpixel accuracy [27]. While both cells maintain a similar biconcave geometry, only the metabolically active cell displayed vigorous, dynamic membrane flickering. This difference was quantitatively confirmed by analyzing the time series of radial displacements (Figure 1B). The live RBC exhibited high-amplitude fluctuations with broad, non-Gaussian displacement distributions, indicative of non-equilibrium active processes. In contrast, the fixed RBC showed dampened, thermally limited motion with narrow Gaussian-like distributions compatible with lacking ATP and active cytoskeletal dynamics. Further insight into the membrane dynamics was obtained by mapping the phase space (displacement vs. velocity) of local fluctuations (Figure 1C). Healthy cells showed extended, elliptical phase portraits characteristic of active, viscoelastic relaxation. Conversely, fixed cells exhibited tightly clustered distributions near the origin, consistent with equilibrium thermal noise and mechanical stiffening. From these datasets, we extracted key mechanical parameters including instantaneous rigidity and effective fluidity of the RBC membrane using the Mason–Weitz microrheological formalism (see Methods). Our analysis confirms that ATP depletion via chemical fixation drastically increases membrane stiffness and reduces cytoskeletal-driven flexibility. These changes are mechanistically tied to the inactivation of metabolic pathways, in particular glycolysis, which fuels the dynamic remodeling of the spectrin–actin cytoskeleton [23,26]. As illustrated in Figure 1D, our analytic methodology based on reporting flickering profiles provides a quantitative mechanical fingerprint of the RBC membrane state [29,32]. Importantly, we here enable functional mechanical comparisons across physiological and pathological RBC populations, including those subjected to metabolic inhibition, parasitic infection, or genetic enzymopathies such as PKLR deficiency. The proposed approach thus offers a powerful tool to map membrane mechanics as a diagnostic proxy of RBC vitality and metabolic fitness in health and disease.

**Figure 1.**
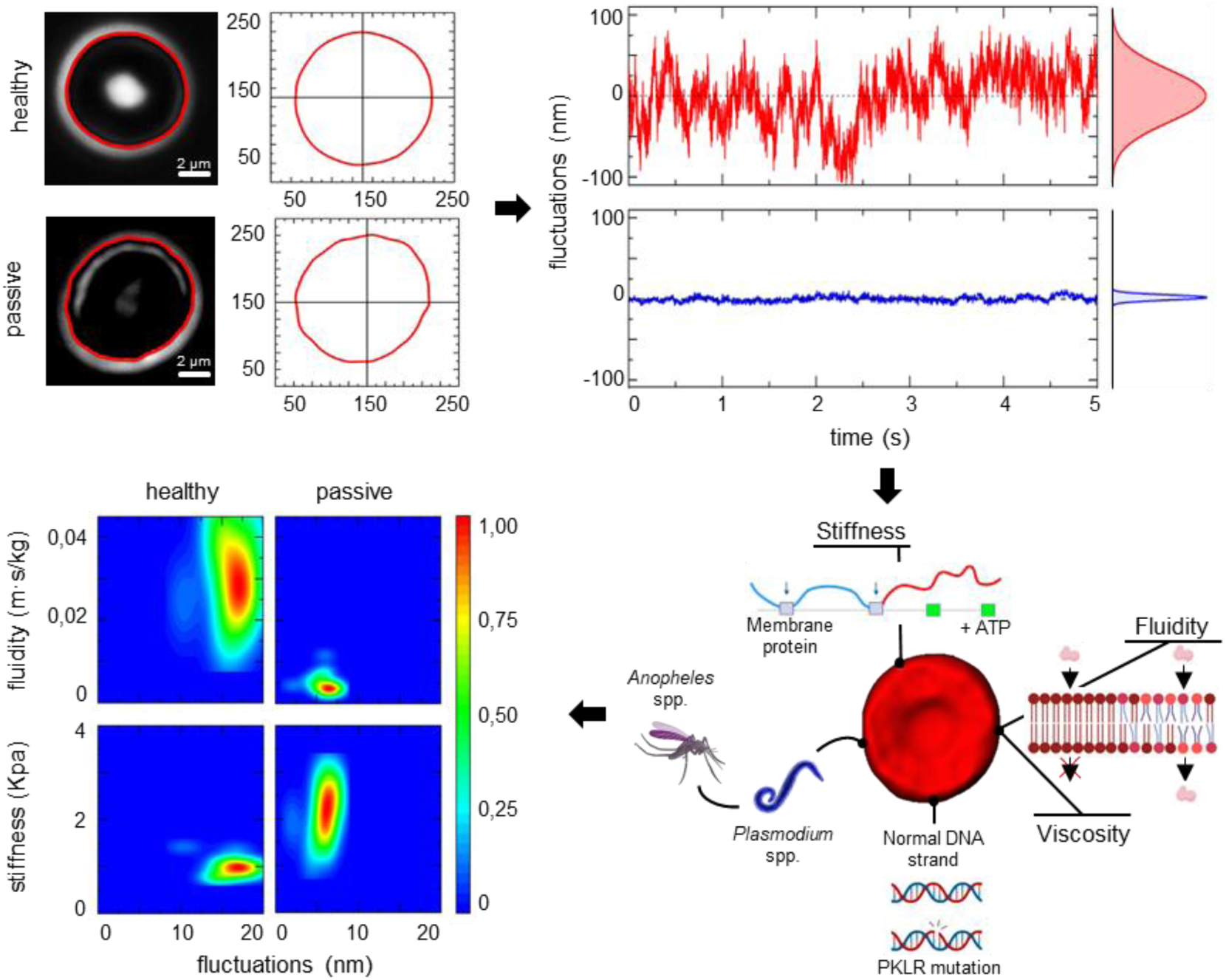
ATP-dependent membrane flickering distinguishes metabolically active from fixed red blood cells (RBCs) from PKLR-deficient C57BL/6J mice. **A)** Phase-contrast micrographs (left) of a live healthy RBC (top) and a glutaraldehyde-fixed RBC (bottom), with their respective equatorial membrane contours overlaid in red. The tracked membrane profiles (center) are circularized and superimposed for morphological comparison. **B)** Representative time series of radial membrane displacements (left) from a tracked point on each cell, and their corresponding probability distributions (right). The live cell (top, red) shows larger-amplitude and more dynamic fluctuations than the fixed cell (bottom, blue), whose motion is thermally constrained. **C)** Two-dimensional phase space distributions of tracked membrane displacements (radial position vs. time derivative) for each condition reveal broader, non-Gaussian excursions in healthy cells (top), characteristic of active flickering, versus restricted, near-thermal behavior in the fixed case (bottom). **D)** Schematic illustrating how RBC membrane flickering emerges from ATP-dependent cytoskeletal activity and is compromised in pathological or fixed conditions. Perturbations such as Plasmodium infection or PKLR enzymopathies alter membrane mechanics through metabolic dysfunction, reflected in altered flickering dynamics.

### Pyruvate kinase deficiency alters the mechanical phenotype of red blood cells under *Plasmodium* infection

To evaluate the impact of PKLR deficiency and malaria infection on RBC membrane mechanics, we performed a comparative biophysical analysis on four mouse-derived RBC populations from PKLR-deficient C57BL/6J mice: (i) healthy wild-type RBCs (hRBC), (ii) wild-type RBCs infected with *Plasmodium yoelii* (iRBC), (iii) uninfected RBCs from PKLR- deficient mice (hPKLR), and (iv) PKLR-deficient RBCs infected with *P. yoelii* (iPKLR). Figure 2 shows experimental results for a total of 60 single-cell specimens per condition analysed by high-speed fluctuation spectroscopy, allowing quantification of local flickering amplitude, fluidity, and stiffness at submicron spatial resolution. Representative phase-contrast images and corresponding microrheological maps of each cell population are shown in Figure 2A. Healthy RBCs displayed highly dynamic membrane fluctuations with spatially heterogeneous distributions of fluidity and moderate stiffness. In contrast, PKLR- deficient cells exhibited overall reduced fluctuation amplitudes and increased mechanical stiffness, consistent with impaired glycolytic activity and altered cytoskeletal remodeling. Notably, *Plasmodium*-infected RBCs (iRBC) retained moderate levels of fluctuation and mechanical softness, although with somewhat reduced heterogeneity relative to hRBCs. Infected PKLR-deficient RBCs (iPKLR) showed the most pronounced mechanical impairment, with markedly dampened fluctuations and consistently high stiffness across the entire cell perimeter. These trends were corroborated quantitatively by statistical boxplot comparisons of average flickering amplitude and stiffness for each population (Figure 2B). Healthy RBCs showed the largest mean fluctuation amplitudes and lowest stiffness values. PKLR deficiency significantly reduced flickering and increased stiffness (p < 0.01), while infection alone induced a moderate but statistically significant softening (p < 0.05). The most rigid and fluctuation-suppressed phenotype corresponded to the double-hit condition (iPKLR), combining enzymopathy and infection. To visualize population-level differences, phase space density plots were constructed for each subpopulation, correlating fluctuation amplitude with fluidity (top row) and stiffness (bottom row) (Figure 2C). Healthy cells were distributed across a wide dynamic range, occupying high-fluctuation, high-fluidity regions. Infected cells (iRBC) clustered in a slightly narrowed distribution but still retained moderate mechanical activity. In contrast, PKLR-deficient cells, both infected and uninfected, were confined to low-fluidity, high-stiffness regions, with minimal spatial dispersion. Together, these findings reveal that PKLR deficiency induces a consistent mechanical rigidification of the RBC membrane, while *Plasmodium* infection exerts more variable effects, potentially reflecting stage-dependent metabolic crosstalk between the parasite and host cell. The observed mechanical profiles distinguish pathological cell states and provide a biophysical fingerprint of metabolic impairment in circulating erythrocytes.

**Figure 2.**
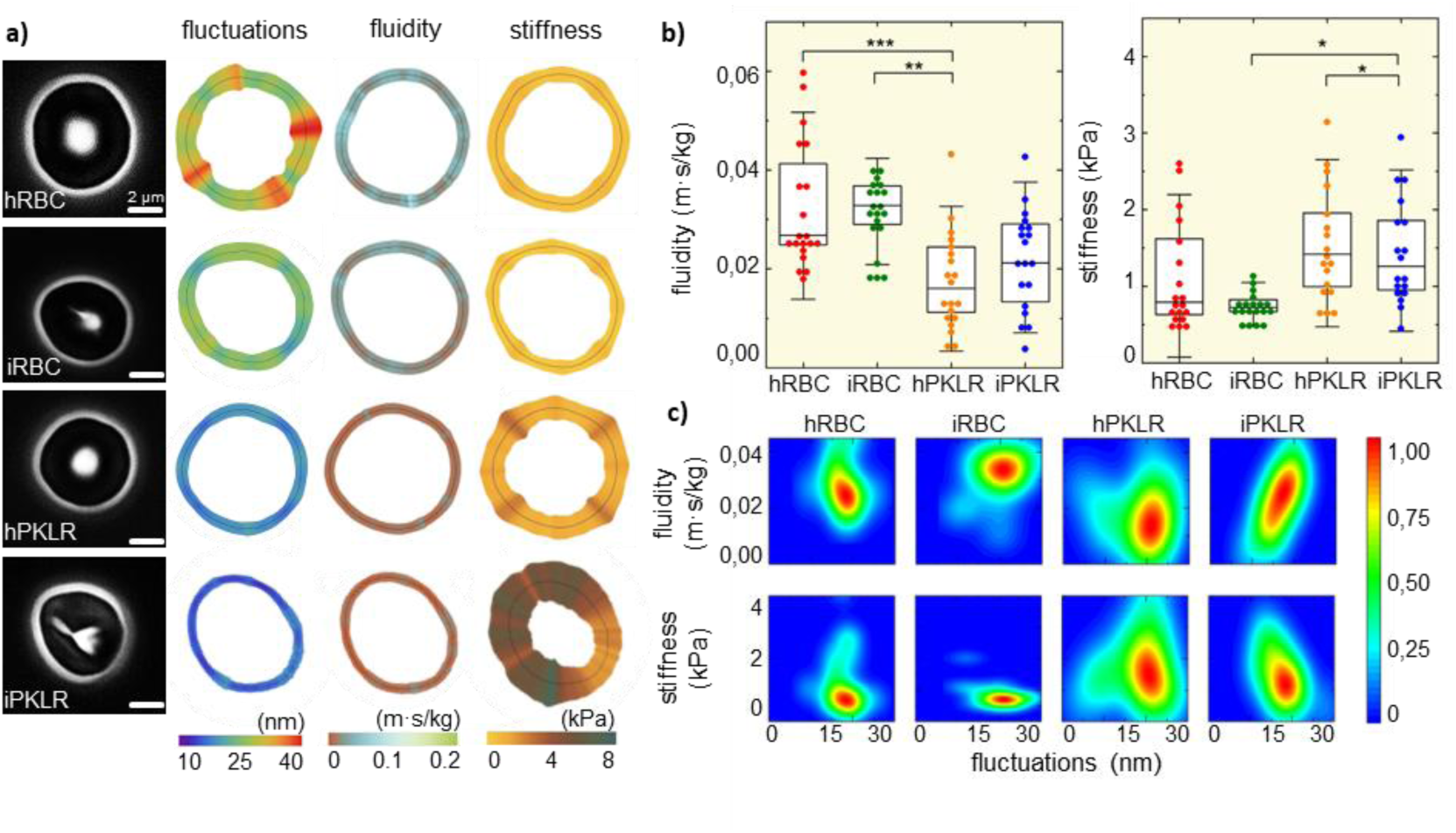
Mechanical phenotype profiling of healthy, infected, and PKLR-deficient red blood cells (RBCs) from PKLR-deficient C57BL/6J mice. **A)** Representative phase-contrast images and quantitative mechanical maps for four RBC populations: healthy RBCs (hRBC), malaria-infected RBCs (iRBC), healthy RBCs from a PKLR-deficient mouse (hPKLR), and infected PKLR-deficient RBCs (iPKLR). From left to right, each row shows the original micrograph, followed by spatial heatmaps of flickering amplitude (nm), local fluidity (pN·s/µm), and stiffness (kPa) derived via microrheological analysis. Color scales indicate relative intensities within each population. Healthy cells exhibit enhanced membrane undulations, distributed fluidity, and moderate stiffness, whereas PKLR-deficient or infected cells show attenuated fluctuations and increased mechanical rigidity. **B)** Boxplots of globally averaged flickering amplitude (left) and stiffness (right) for the four populations. iPKLR cells exhibit the lowest flickering activity and highest stiffness, while hRBCs display the highest flickering and softest mechanics. Statistical significance determined by ANOVA with post-hoc Tukey correction (*p < 0.05, **p < 0.01, ***p < 0.001). **C)** Phase space maps displaying joint distributions of flickering amplitude versus fluidity (top row) and versus stiffness (bottom row) across all single-cell measurements. Kernel density estimates highlight characteristic mechanical signatures of each RBC population. Healthy cells cluster in regions of high fluctuation and high fluidity, while PKLR-deficient and infected cells are progressively shifted toward stiffer and less dynamic profiles. These results reveal distinct mechanical phenotypes associated with infection and glycolytic impairment, offering a potential diagnostic readout for RBC vitality and pathology.

### Metabolic Inhibition Progressively Suppresses Flickering and Mechanical Fluidity in Healthy Red Blood Cells

To assess the influence of metabolic activity on RBC membrane mechanics, we compared healthy RBCs (hRBC) to two metabolically impaired states from wild-type C57BL/6J mouse strain: ATP- depleted cells treated with iodoacetamide and inosine (“drugged” RBCs) and fully fixed cells treated with glutaraldehyde (“passive” RBCs). High-speed phase-contrast microscopy combined with passive microrheology analysis was used to quantify flickering dynamics, fluidity, and stiffness in each condition. As shown in Figure 3A, healthy RBCs displayed heterogeneous membrane fluctuations and relatively soft mechanical behavior, with spatially distributed areas of enhanced fluidity. In contrast, drugged cells exhibited visibly reduced fluctuation amplitudes and localized stiffening, while fixed cells presented uniformly suppressed dynamics and elevated stiffness. Quantitative comparison (Figure 3B) revealed a significant decrease in membrane fluctuation amplitude and a concurrent increase in stiffness between healthy, drugged, and passive cells (n = 20 per group; ANOVA with post hoc testing, ***p < 0.001). This confirms that both partial and total metabolic inhibition translate into progressive mechanical rigidification. Finally, phase-space density plots (Figure 3C) demonstrated a shift in population distributions: healthy cells occupied a broad domain characterized by high fluidity and moderate stiffness; drug-treated RBCs shifted toward lower fluidity and higher stiffness; and fixed cells clustered tightly in a mechanically inactive regime. These findings corroborate the ATP-dependence of RBC flickering and highlight the sensitivity of membrane viscoelasticity to metabolic impairment.

**Figure 3.**
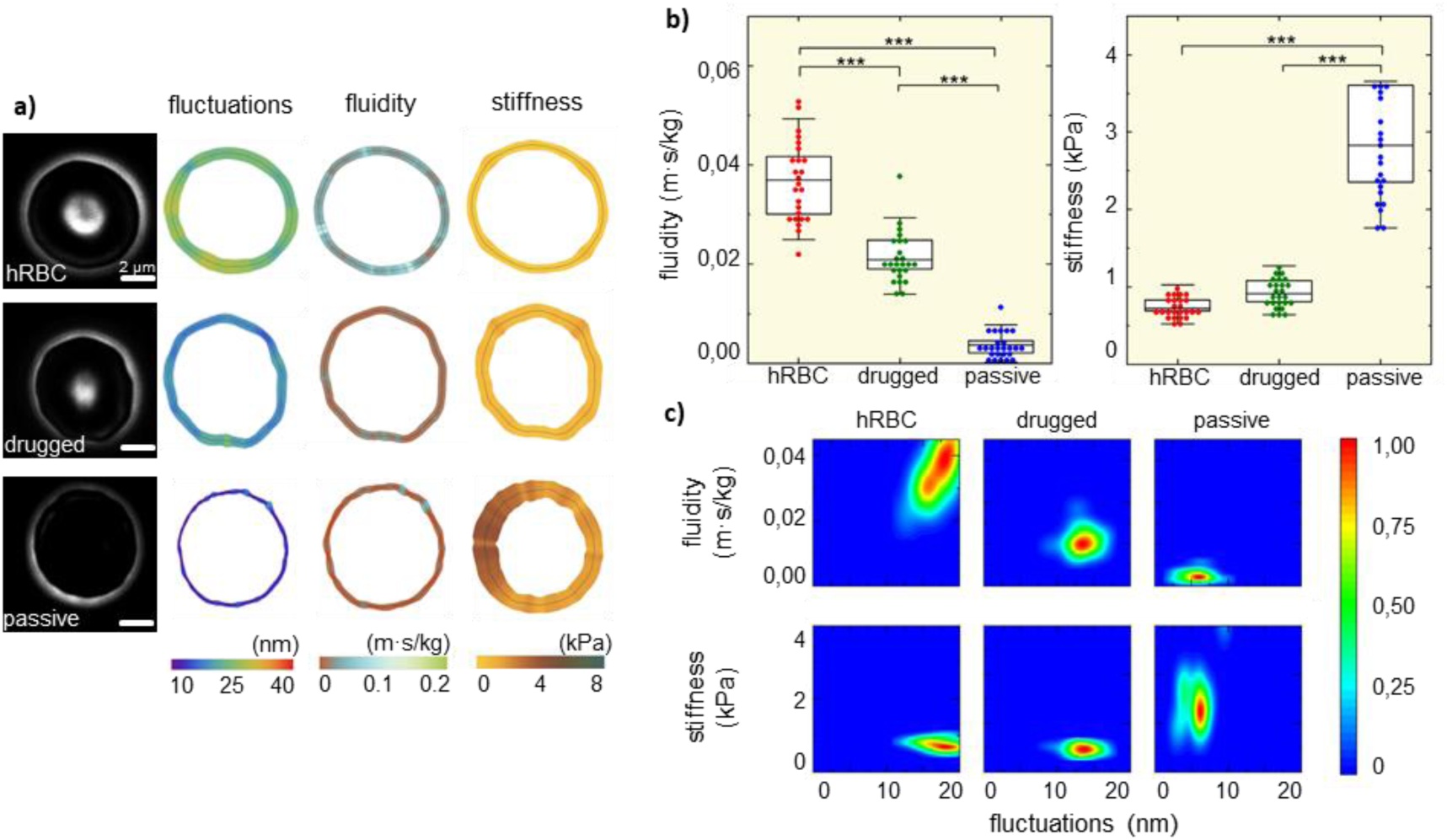
Metabolic inhibition selectively alters red blood cell membrane mechanics. Healthy red blood cells (hRBCs) from wild-type C57BL/6J mice were subjected to two conditions of metabolic impairment: drug-induced ATP depletion (“drugged”) and complete chemical fixation (“passive”), and compared to untreated controls. **A)** Representative phase-contrast micrographs (left column) show the circular morphology of hRBCs across conditions. Spatially resolved microrheological maps reveal the distributions of (second column) flickering amplitude, (third column) fluidity, and (fourth column) stiffness across the membrane contour. Healthy cells display heterogeneous fluctuations and mechanical softening, whereas ATP-depleted (“drugged”) and fixed (“passive”) RBCs exhibit marked reductions in fluctuation and fluidity, with increased rigidity. Color gradients reflect the normalized scales of each parameter for comparative visualization. **B)** Quantitative comparison of flickering amplitude (left) and stiffness (right) across the three groups (n = 20 per condition). Boxplots indicate median (central bar), interquartile range (box), and full data spread (whiskers and outliers). Statistically significant differences were found between all groups (***p < 0.001, ANOVA with post hoc test), demonstrating that both metabolic inhibition and chemical fixation significantly reduce dynamic membrane behavior and increase stiffness. **C)** Phase-space density maps illustrate the population-level distributions of fluidity and stiffness as a function of flickering amplitude. Healthy RBCs occupy a broad region of high fluctuations and fluidity, while drug-treated cells are more constrained and centered around lower values. Fixed cells collapse into narrow, rigid, and fluctuation-suppressed states, indicative of fully passive, non-metabolic behavior. These results confirm that RBC flickering activity is highly dependent on metabolic state and ATP availability, and that progressive energetic inhibition leads to a biophysically distinguishable loss of mechanical adaptability.

## DISCUSSION

Red blood cell (RBC) deformability is essential for microcirculatory (patho)physiology [1–4,6]. RBC mechanical properties rely on the fine-tuned interplay between the lipid bilayer and the underlying spectrin-based cytoskeleton [14,15], a relationship that is actively regulated by ATP-dependent processes [4]. In this study, we employed label-free flickering spectroscopy, combined with passive microrheology, to characterize how genetic and parasitic challenges affect the mechanical state of RBC membranes. Our biophysical results demonstrate that intracellular ATP availability—regulated primarily through anaerobic glycolysis, is a key determinant of RBC viscoelasticity and active membrane behavior. To define the limits of metabolically driven RBC mechanics, we first established a baseline by comparing active healthy cells to glutaraldehyde-fixed passive controls. Fixed cells, lacking metabolic activity, exhibited a dramatic suppression of flickering and a substantial increase in membrane stiffness. These passive cells represent a mechanical “ground state”, useful for benchmarking metabolically active fluctuations. This observation confirms previous reports that ATP is required to sustain flickering dynamics [23–33], and supports the use of high-speed flickering analysis as a sensitive, non-invasive indicator of RBC energetic status [27].

Hence, we next assessed the effects of two biologically relevant stressors: infection with *Plasmodium yoelii* (a murine model of malaria), and genetic pyruvate kinase deficiency (PKD) caused by *PKLR* mutation. RBCs infected with *P. yoelii* in the early sporozoite stage retained moderate flickering and fluidity, indicating that early parasite development does not immediately deplete host ATP reserves. This finding is in line with previous studies showing that Plasmodium co-opts host energy metabolism during the early stages of infection [44]. By contrast, *PKLR*-deficient RBCs —whether infected or not, exhibited markedly reduced flickering amplitudes and increased stiffness, consistent with impaired ATP production through the Embden–Meyerhof glycolytic pathway. These mechanical changes mirror the clinical phenotype of PKD patients, who present with rigid, osmotically fragile RBCs and chronic hemolytic anemia [6,7]. Additionally, pharmacological ATP depletion using iodoacetamide and inosine recapitulated this rigidification phenotype, reinforcing the direct link between metabolic energy availability and cytoskeletal softening [23,26,33].

### PKLR Deficiency as a Rigidity Barrier to Malaria Infection

A significant finding of this study is that *PKLR*-deficient RBCs, when infected with *Plasmodium yoelii*, retain the mechanical characteristics of ATP-depleted cells—namely, reduced flickering and increased stiffness, even in the presence of intracellular parasites. This suggests that parasite-induced remodeling of the host membrane is either delayed or functionally impaired in these metabolically compromised cells. We therefore hypothesize that *PKLR*-deficient RBCs act as a **mechanical barrier to malaria progression**, where the rigidity of the membrane, stemming from insufficient ATP to support cytoskeletal rearrangements, impairs the parasite’s ability to modify the host cell, thereby obstructing its maturation beyond the early sporozoite stage. This form of resistance is distinct from immune-mediated responses and instead reflects non-immunological, mechanically driven incompatibility rooted in altered host metabolism. Supporting this hypothesis, we observed that infected PKLR-deficient cells lacked the typical spectrum of membrane remodelling seen in metabolically active infected RBCs.

Although further validation is required, our findings resonate with a growing body of evidence suggesting that biomechanical membrane host incompatibility can act as a constraint on *Plasmodium* invasion [50]. For instance, computational modelling by Alimohamadi et al. (2023) demonstrated that increased membrane tension raises the energetic barrier for merozoite wrapping and entry, thereby reducing invasion efficiency [39]. Similarly, the Dantu blood group variant, associated with increased membrane stiffness, has been shown to confer partial protection against severe malaria by limiting parasite access to the cytosol, as reported by Kariuki et al. (2020) [38]. In both cases, altered mechanical properties of the host membrane restrict parasite invasion and development.

Our results suggest that PKLR-linked metabolic stiffening may produce a comparable phenotype, thus providing a novel perspective on how host metabolic disorders can contribute to innate resistance against malaria. These findings are further supported by in vitro studies demonstrating that elevated levels of 2,3-diphosphoglycerate—a metabolic hallmark of PKLR deficiency—not only impair *Plasmodium falciparum* development [51] but also alter RBC mechanics in ways that may hinder parasite invasion [11]. Together, these results reinforce the concept that metabolic and biomechanical changes in PKLR-deficient RBCs synergize to form a physical barrier to parasite progression.

### Cross-Species Evidence Supporting PKLR Deficiency as a Biomechanical Defense Against Malaria

Our findings in *PKLR*-deficient C57BL/6J mice support the broader concept that metabolically induced rigidity of red blood cells (RBCs) may serve as a conserved protective mechanism against malaria infection. Notably, this aligns with previous studies that have explored the relationship between *PKLR* deficiency and resistance to malaria across multiple experimental models. A seminal study by Min-Oo et al. (2007) demonstrated that the severity of pyruvate kinase (PK) deficiency correlates with enhanced survival following *Plasmodium chabaudi* infection in different murine genetic backgrounds, including CBA and BALB/c mice [46]. This work highlighted that host erythroid metabolic status could impair parasite replication in vivo, specifically, a reduced ATP production due to *PKLR* mutation.

Our results extend these findings to the C57BL/6J strain, underscoring the robustness of this protective phenotype across mouse lineages. Moreover, in vitro studies in human RBCs have provided further validation. For example, research by Ayi et al. (2004) showed that *P. falciparum* invasion is significantly reduced in human erythrocytes harboring *PKLR* mutations, suggesting that metabolic stiffening can confer resistance across species boundaries [52]. These findings complement our current data showing that RBCs from *PKLR*-deficient mice maintain a rigid mechanical phenotype —even upon *Plasmodium yoelii* infection, thereby supporting a model in which ATP-dependent flickering and deformability are necessary for successful parasitic remodeling. Taken together, these cross-species studies converge on a unifying wisdom: *genetically impaired glycolysis in RBCs generates a mechanically resistant host environment that limits malaria progression*.

Our high-resolution analysis of flickering amplitude and membrane stiffness now provides a biophysical explanation for these observations, revealing that reduced ATP availability suppresses the active, flickering-based mechanical plasticity that parasites exploit for invasion and survival. Finally, these insights also raise broader questions about genotype–phenotype variability in *PKLR* deficiency. Human populations exhibit a spectrum of mutations in the *PKLR* gene, with varying degrees of enzymatic activity, clinical expression, and potential protective effects against malaria. Beyond host-derived membrane mechanics, emerging evidence also implicates parasite effectors in modulating RBC biomechanics. *P. yoelii*, for instance, employs the erythrocyte-binding ligand PyEBL to mediate host cell entry. Notably, mutations in PyEBL have been shown to alter the mechanical and immune landscape of infected RBCs, including enhanced membrane fragility and phosphatidylserine exposure, without impairing invasion efficiency [53]. This supports a broader view: membrane remodeling—whether host-imposed or parasite-induced—is a critical determinant of infection outcome, independent of invasion per se.

Future translational studies combining genetic, mechanical, and infection-based profiling may uncover how this phenotypic diversity shapes host–parasite dynamics and may inform strategies for personalized risk assessment or therapeutics based on RBC biomechanical phenotyping. Finally, while single-cell measurements revealed statistically significant mechanical differences across conditions, we recognize the potential influence of inter-animal variability. To address this, future analyses will incorporate mixed-effects models and expanded biological replicates to better disentangle intra- from inter-individual sources of variance. Beyond this proof-of-concept study, follow-up preclinical work will leverage larger animal cohorts and hierarchical statistical frameworks to rigorously quantify variability across genetic and environmental contexts and ensure the robustness and generalizability of the observed mechanical phenotypes.

## CONCLUSIONS AND OUTLOOK

This study demonstrates that red blood cell (RBC) flickering is a sensitive, ATP-dependent mechanical phenotype that reflects the energetic and structural integrity of the cell. By combining high-speed phase-contrast imaging with passive microrheology, we established a powerful, label-free approach to quantify RBC membrane mechanics across genetic (PKLR mutation), metabolic, and parasitic challenges. We show that flickering is not merely a passive thermal artifact but an active, energy-sustained viscoelastic process. In ATP-depleted conditions, whether genetically (PKLR deficiency), chemically, or pathogenically induced, the RBC membrane becomes mechanically rigid and flickering collapses. This dynamic stiffening constrains membrane plasticity, which is crucial for the *Plasmodium* life cycle. In PKLR-deficient RBCs, the resulting rigidity likely hampers parasite remodeling capacity, offering a non-immunological, mechanical barrier to infection. This finding positions PKLR- linked enzymopathy not only as a marker of hematologic dysfunction but as a functional constraint on parasite development, parallel to the protective effects observed in other mechanical phenotypes such as the Dantu variant or sickle trait. Our biophysical results on RBC flickering align with previous biochemical studies showing that increased RBC tension reduces invasion efficiency by elevating the energy cost of merozoite membrane wrapping.

Our findings elevate RBC flickering as a functional biomarker and potential mechanical defense mechanism, shaped by the interplay between bioenergetics, membrane architecture, and pathogen adaptation. Future work should examine how different parasite stages and ligand variants (e.g., PyEBL or PfRH families) interact with varying host biomechanical states, including PKLR heterozygosity in humans. As a quantitative diagnostic tool, flickering spectroscopy offers a scalable, non-invasive platform to assess RBC metabolic health, pathogen susceptibility, and therapeutic responses. Integrating this technique with molecular profiling (e.g., invasion ligands, cytoskeletal regulators, and phospholipid asymmetry) may unlock multimodal biomarkers for precision hematology and malaria management. Beyond its relevance in murine models, our findings offer translational value for understanding human PK deficiency and its implications for malaria susceptibility. Given the clinical diversity of PKLR mutations in humans—ranging from asymptomatic carriers to severe hemolytic phenotypes—membrane flickering analysis may provide a functional stratifier of disease severity and infection risk. Integrating flickering-based nanomechanical profiling into existing diagnostic platforms, such as laser ektacytometry or flow cytometry, could enable non-invasive assessment of RBC deformability and metabolic competence, supporting personalized risk assessment in both hereditary anemia and malaria-endemic settings.

## Acknowledgments

NH is contracted by INVESTIGO-CM Program from Comunidad de Madrid (Grant number 09-PIN1-00009.8/2022). LHM is contracted by María Zambrano Program from Ministerio de Universidades de España for the attraction of international talent under Next Generation European Union funding (grant CT19/22). The work was supported by the Spanish Ministry of Science and Innovation (MICINN– Agencia Española de Investigación AEI) under grants PID2019-108391RB-100 and TED2021-132296B-C52 (to FM), PID2023-149637OB-I00 (to JMB), PID2023-152564OB-I00 and PLEC2023-010243 (to JCS); and Comunidad de Madrid under grants S2018/NMT-4389 and Y2018/BIO-5207 (to FM). This study was also funded by the REACT-EU program PR38-21-28 ANTICIPA-CM, a grant by Comunidad de Madrid and European Union under FEDER program, from European Union in response to COVID-19 pandemics. The funders had no role in the study design, data collection, analysis, preparation of the manuscript, or the decision to publish.

